# AAV9 gene therapy restores lifespan and treats pathological and behavioral abnormalities in a mouse model of CLN8-Batten disease

**DOI:** 10.1101/2020.05.05.079350

**Authors:** Tyler B. Johnson, Katherine A. White, Jacob T. Cain, Logan Langin, Melissa A. Pratt, Jon Brudvig, Clarissa D. Booth, Derek J. Timm, Samantha S. Davis, Brandon Meyerink, Shibi Likhite, Kathrin Meyer, Jill M. Weimer

**Affiliations:** Pediatrics and Rare Diseases Group, Sanford Research, Sioux Falls, SD USA; Amicus Therapeutics, Philadelphia, PA USA; The Research Institute at Nationwide Children’s Hospital, Columbus, Ohio, USA; The Department of Pediatrics, The Ohio State University, Columbus, Ohio, USA

## Abstract

CLN8 disease is a rare form of neuronal ceroid lipofuscinosis caused by biallelic mutations in the *CLN8* gene, which encodes a transmembrane endoplasmic reticulum protein involved in trafficking of lysosomal enzymes. CLN8 disease patients present with myoclonus, tonic-clonic seizures, and progressive declines in cognitive and motor function, with many cases resulting in premature death early in life. There are currently no treatments that can cure the disease or substantially slow disease progression. Using a mouse model of *CLN8* disease, we tested the safety and efficacy of an intracerebroventricularly (ICV)-delivered self-complementary AAV9 (scAAV9) gene therapy vector driving expression of human *CLN8*. A single neonatal injection was safe and well-tolerated, resulting in robust transgene expression throughout the brain and spinal cord from 4 to 24 months, reducing histopathological and behavioral hallmarks of the disease and completely restoring lifespan from 10 months in untreated animals to beyond 24 months of age in treated animals. These results demonstrate, by far, the most successful rescue reported in an animal model of CLN8 disease, and supports gene therapy as a promising therapeutic strategy for this disorder.

## Introduction

The neuronal ceroid lipofuscinoses (NCLs, also known as Batten Disease) are a phenotypically similar group of rare lysosomal storage disorders caused by mutations in one of at least 13 identified ceroid-lipofuscinosis neuronal (CLN)-related genes^1^, and are collectively the most prevalent neurodegenerative disease in children (incidence 2-4/100,000 births)^2, 3^. Patients typically present with CNS-related symptoms including mental and motor deficits, blindness, and epilepsy in the first decade of life, with variable symptomatology, onset, and progression depending on the genetic subtype. All forms of NCL are progressive, most are fatal, and none yet have a cure^1^.

At the cellular level, NCLs are characterized by the lysosomal accumulation of autofluorescent storage material (ASM) consisting of lipid-based lipofuscin and other waste products^4^. Some CLN genes encode soluble lysosomal enzymes and are thus linked directly to lysosomal activity (CLN1, CLN2, CLN10, CLN13), while others have more elusive function^1^. Recently, CLN8, a transmembrane endoplasmic reticulum (ER) protein, was shown to regulate the ER to Golgi trafficking of catabolic enzymes destined for the lysosome^5^. In the absence of CLN8, lysosomal enzymes are depleted, leading to classic NCL CNS pathology including ASM accumulation, glial activation, and neurodegeneration throughout the CNS^6^.

Mutations in *CLN8* can lead to an aggressive form of variant late infantile NCL in human patients that typically presents in childhood with myoclonus, tonic-clonic seizures, and progressive motor decline and dementia. Visual deficits or blindness can develop throughout the disease course^7-12^, or not at all^12-14^. Phenotypic severity varies from milder forms, such as the “Northern Epilepsy” caused by missense mutations in the Finnish population^12^, to more severe forms caused by different sets of mutations, including null alleles^15, 16^. Various small molecule treatments have shown some efficacy in animal models of CLN8 disease^17-19^, yet none have been adopted in the clinic, resulting in an unmet need to halt or reverse disease progression.

Recently, gene therapy has shown promise for treating a variety of genetic disorders, including the NCLs. Adeno-associated virus (AAV)-based therapies have shown some of the greatest potential, with low immunogenicity and toxicity, persistent gene expression, and favorable tropism patterns^20^. AAV-based therapies have been tested in animal models for CLN1, CLN2, CLN3, CLN6, and CLN10, and are in clinical trials for human CLN2, CLN3, and CLN6 patients^1^. While AAV-based therapies have been shown to be generally safe and well-tolerated, they have not yet been explored for CLN8 disease. Here, we use the *Cln8*^*mnd*^ mouse model to test the efficacy of AAV serotype 9 (AAV9) gene therapy for CLN8 disease.

The Cln8 motor neuron degeneration (*Cln8*^*mnd*^) mouse, a naturally occurring mutant that has been extensively characterized^21^, exhibits a disease pattern with strong similarity to human *CLN8*^*mnd*^ disease patients. Cln8 homozygotes display early and progressive neuroinflammation and motor neuron degeneration culminating in paralysis and early death around one year of age^21-23^. The causative mutation is a single base pair insertion (267-268insC) resulting in a frameshift and premature stop codon^12^. Microglial activation is apparent at the early symptomatic stage of 3 months and is followed shortly thereafter by progressive astrocytosis and thalamic and cortical neurodegeneration beginning at 5 months of age^6^. Associative learning deficits are detectable as early as 14-16 weeks of age, along with reductions in electroretinogram (ERG) responses and pupillary light reflexes^24^. Retinal photoreceptor death begins as early as postnatal day 21 (P21)^25^.

We utilized an intracerebroventricular (ICV)-delivered self-complementary AAV9 (scAAV9) gene therapy vector, based on our prior success with this treatment in a mouse model of CLN6 disease^26^. Our vector (scAAV9.pT-MecP2.CLN8) utilizes an AAV9 capsid combined with truncated *MecP2* promoter (pT-MecP2).This combination achieves widespread and persistent transduction throughout the CNS, including in the cortex and thalamus, following a single injection. We quantified outcomes by measuring well characterized disease correlates including cortical and thalamic lysosomal storage and glial activation, behavioral assays of motor performance and learning/memory, and survival.

## Results

### scAAV9.pT-MecP2.CLN8 drives widespread expression of hCLN8 through 24 months of age in Cln8^mnd^ mice

To express full length human *CLN8* (hCLN8) in the CNS of treated animals, we designed an AAV9 viral vector expressing *hCLN8* under the control of the 546 promoter: scAAV9.pT-MecP2.CLN8. The AAV9 capsid has been shown to be translatable between animal models of neurodegenerative disease and human subjects^27, 28^, and is currently being used in clinical trials for CLN3 and CLN6 disease (Clinicaltrials.gov Identifiers: NCT03770572 and NCT02725580, respectively) following successful preclinical studies^26, 29^. scAAV9.pT-MecP2.CLN8 drives expression with a truncated version of the *MecP2* promoter (pT-MecP2), which has been shown to achieve physiologically appropriate expression levels for genes that are endogenously expressed with low abundance^29^. For this study, we delivered scAAV9.pT-MecP2.CLN8 via a single ICV injection at postnatal day one (P1) in male and female *Cln8*^*mnd*^ mice at a dose of 5 x 10^10^ vg (the same dose used in our preclinical CLN6 studies)^26^. PBS was delivered to *Cln8*^*mnd*^ controls.

To characterize spatiotemporal patterns of transgene expression, we examined *hCLN8* transcript using RNAscope, a modified *in situ* hybridization assay. *hCLN8* transcript was present in scAAV9.pT-MecP2.CLN8-treated *Cln8*^*mnd*^ mice in all brain regions examined from 4-24 months, with the most robust levels in the cerebral cortex (motor, somatosensory, and visual cortices), and lower levels in the thalamus and hindbrain (Figure 1). Transcript levels and regional patterns were sustained over the 24-month period. Transcription throughout the somatosensory thalamocortical pathway was of particular interest, as these areas have been shown to be particularly vulnerable in *Cln8*^*mnd*^ mice^6^. scAAV9.pT-MecP2.CLN8 also resulted in transcription of hCLN8 in the cervical, thoracic, and lumbar spinal cord; and in the kidney, as measured by qPCR (Figure 2A-E).

**Figure 1.**
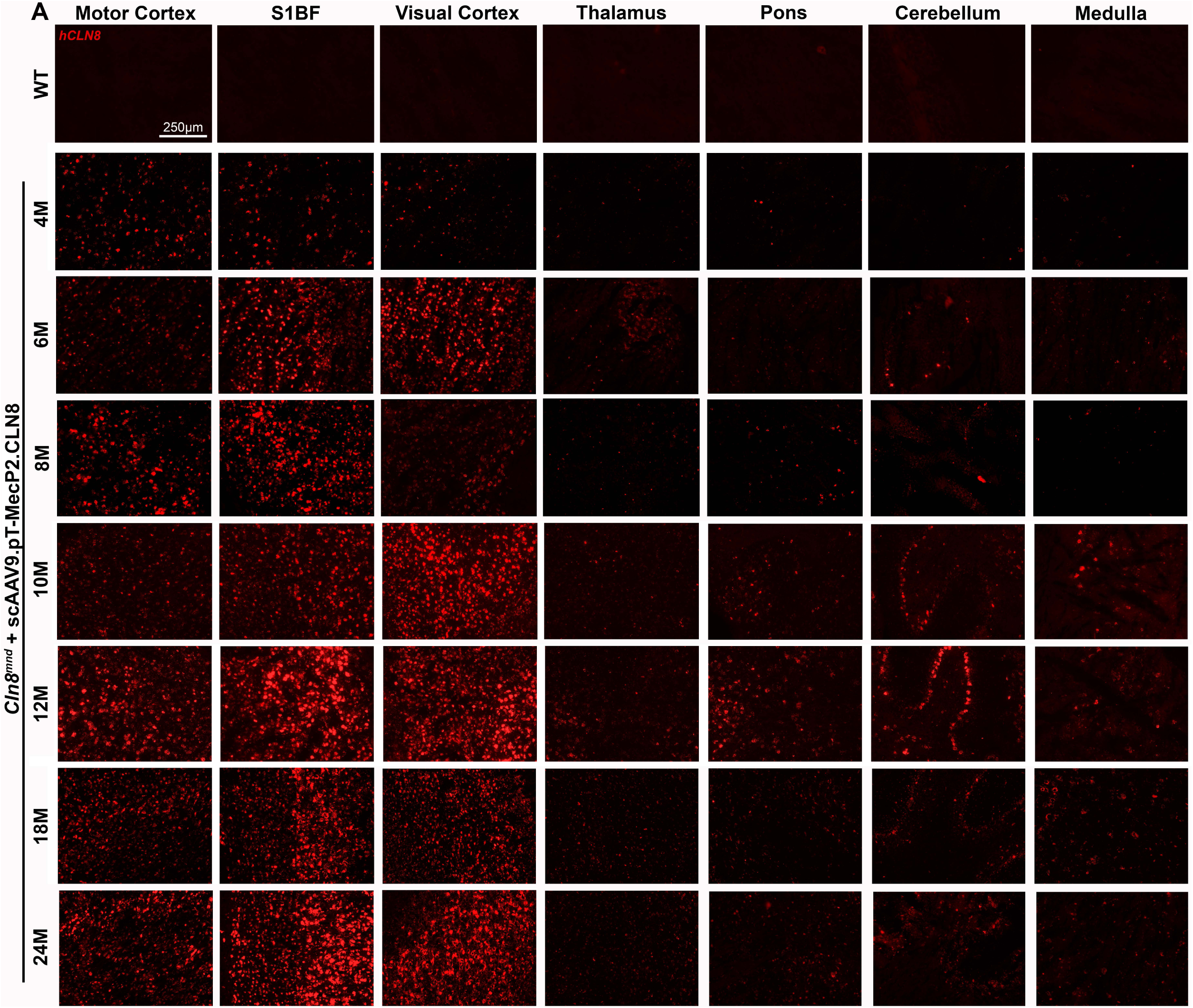
scAAV9.pT-MecP2.CLN8 produces sustained *hCLN8* transcription in the brain of *Cln8*^*mnd*^ mice. A single P1 ICV injection of scAAV9.pT-MecP2.CLN8 results in transcription of *hCLN8* in multiple areas of the brain until 24 months of age (A). The highest transcription levels were observed in the motor cortex, S1BF, and visual cortex, with lower levels of transcription in the thalamus, pons, cerebellum, and medulla.

**Figure 2.**
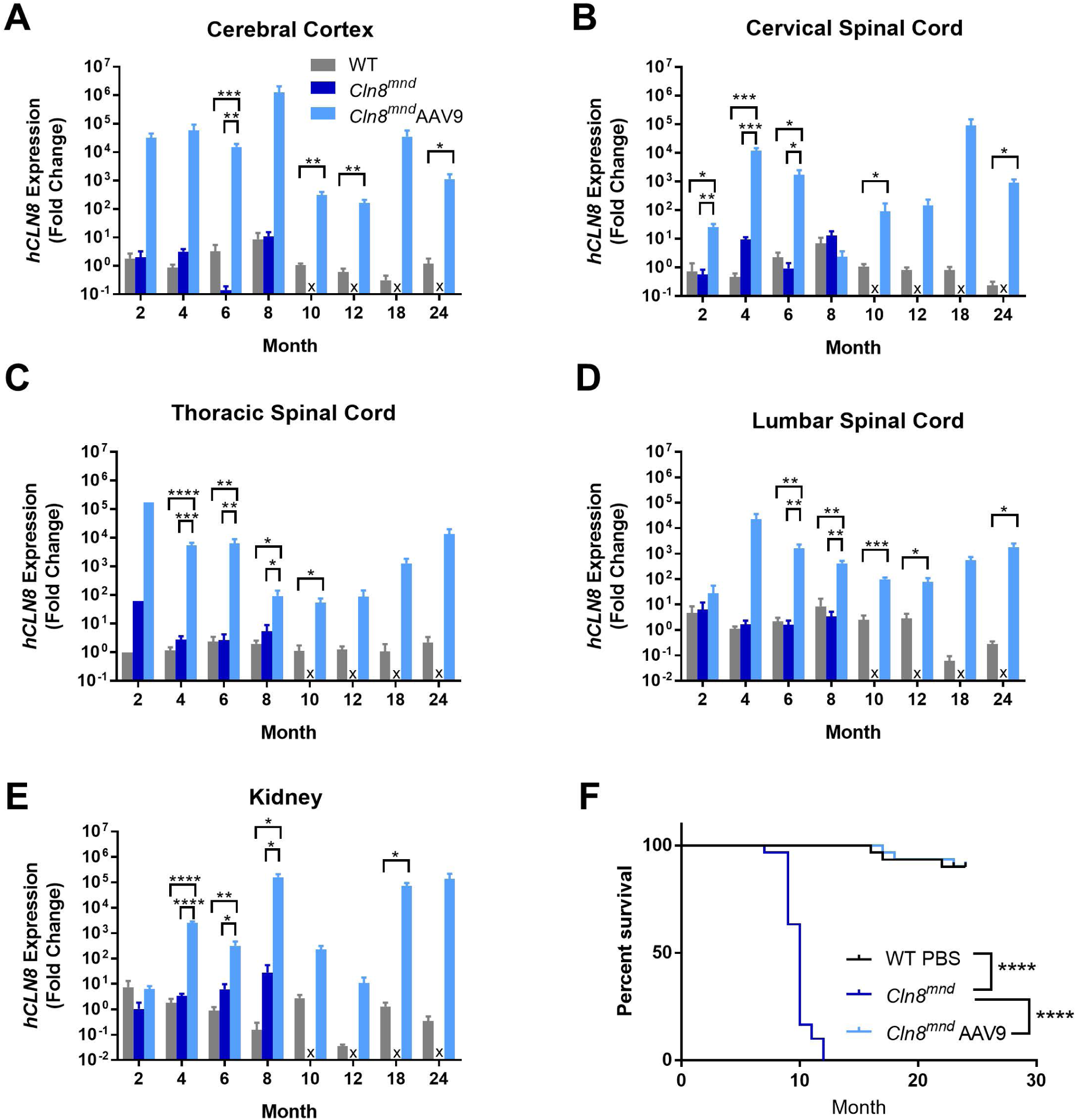
scAAV9.pT-MecP2.CLN8 produces sustained *hCLN8* transcription in the CNS, resulting in enhanced *Cln8*^*mnd*^ survival. As measured by qPCR, P1 ICV administration of scAAV9.pT-MecP2.CLN8 produces increased *hCLN8* transcript in the cerebral cortex (A), cervical spinal cord (B), thoracic spinal cord (C), lumbar spinal cord (D), and kidney (E). Importantly, *hCLN8* expression is sustained throughout the at 24 months post injection. ScAAV9.pT-MecP2.CLN8 extends median lifespan in *Cln8*^*mnd*^ mice from 10 months to beyond 24 months of age (F). Mean ± SEM. Each time point was run independently and analyzed as either a one-way ANOVA, Fisher’s LSD (2 to 8 months), or unpaired t-test (10 to 24 months). Survival curve was analyzed using a Mantel-Cox log-rank test, N=30-31. Detailed sample Ns are described in Supplemental Table 2. *p<0.05, **p<0.01, ***p<0.001, ****p<0.0001.

### scAAV9.pT-MecP2.CLN8 completely rescues lifespan in *Cln8*^*mnd*^ mice

*Cln8*^*mnd*^ mice have a drastically shortened lifespan, with premature death occurring at 7-12 months of age (depending on genetic background), presumably from seizures^23, 30^. In our study, PBS-treated *Cln8*^*mnd*^ mice (C57BL/6J) died between seven and 12 months of age with a median survival of 10 months, while scAAV9.pT-MecP2.CLN8-treated *Cln8*^*mnd*^ mice had a normal lifespan, with a median survival beyond 24 months (Figure 2F). scAAV9.pT-MecP2.CLN8-treated *Cln8*^*mnd*^ mice lived significantly longer than PBS-treated *Cln8*^*mnd*^ mice, and were indistinguishable from wild type (WT) mice in terms of survival curves. This complete rescue of survival far surpasses what was previously the greatest survival increase reported for a therapeutic in *Cln8*^*mnd*^ mice: a 16% increase in mean lifespan with carnitine supplementation^17^.

Additionally, gross necropsy was performed on each mouse at the time of sacrifice. In general, no abnormalities were observed in scAAV9.pT-MecP2.CLN8-treated *Cln8*^*mnd*^ mice in any of the organs examined, including CNS, heart, lung, liver, spleen, kidney, small intestine, and skeletal muscles. Two of the 30 WT animals were euthanized during the study due to labored breathing and mobility issues, and a third WT animal was found dead with an intestinal tumor and internal hemorrhaging (Supplemental Table 1). Of the 31 scAAV9.pT-MecP2.CLN8 treated animals monitored for survival, one animal was euthanized due to a prolapsed rectum, while two animals were euthanized at advanced ages due to abdominal masses that impaired movement. This greatly contrasts the 30 untreated animals that perished by 12 months of age due to seizures, impaired mobility, and/or failure to thrive. Overall, we concluded that the treatment was safe and well-tolerated, with no signs of adverse effects.

### scAAV9.pT-MecP2.CLN8 improves lysosome pathology in *Cln8*^*mnd*^ mice

Intracellular autofluorescent storage material (ASM) accumulates in all forms of NCL, including CLN8, and is frequently used as a correlate for cellular disease burden^23, 31^. While ASM is apparent in many tissues and cell types, it is most evident in neurons^31^. We examined ASM in WT mice and scAAV9.pT-MecP2.CLN8-treated and PBS-treated *Cln8*^*mnd*^ mice from 2 to 24 months. ASM was evident in the ventral posteromedial and ventral posterolateral nuclei of the thalamus (VPM/VPL) and in the primary somatosensory cortex barrel field (S1BF) at two months of age in PBS-treated *Cln8*^*mnd*^ mice, and accumulated rapidly until premature death between eight and 10 months of age (Figure 3A and 3B). scAAV9.pT-MecP2.CLN8 greatly reduced ASM in the VPM/VPL and S1BF at two, four, six, and eight months of age. At four and six months of age, scAAV9.pT-MecP2.CLN8-treated *Cln8*^*mnd*^ mice had significantly more ASM in the VPM/VPL than WT mice, but levels were still far below those of PBS-treated *Cln8*^*mnd*^ mice. At two and eight months of age in the VPM/VPL, and at all time points from two through eight months of age in the S1BF, ASM levels in WT and scAAV9.pT-MecP2.CLN8-treated *Cln8*^*mnd*^ mice were indistinguishable.

**Figure 3.**
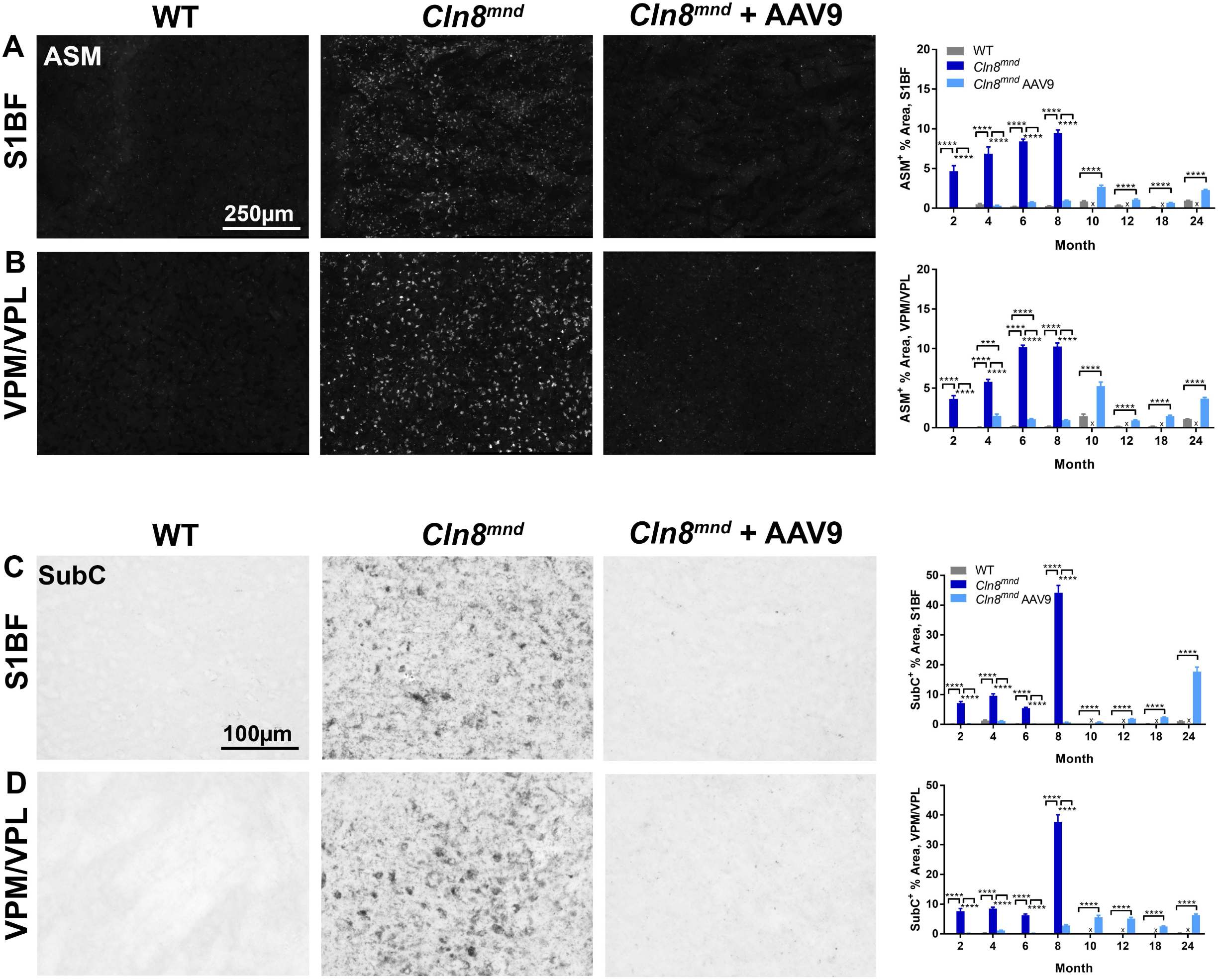
scAAV9.pT-MecP2.CLN8 treatment prevents storage material accumulation in *Cln8*^*mnd*^ mice. As measured by immunohistochemistry, scAAV9.pT-MecP2.CLN8 treatment significantly prevents autofluorescent storage material (ASM; A,B) and mitochondrial ATP synthase subunit c accumulation (SubC; C,D) in the somatosensory cortex (S1BF) and thalamus (VPM/VPL) from two through 24 months of age. Images representative of 8 months of age. Mean ± SEM. Each time point was analyzed independently as either a one-way ANOVA, Fisher’s LSD (2 to 8 months), or unpaired t-test (10 to 24 months). Biological N=4/sex/genotype. Detailed sample Ns are described in Supplemental Table 2. ***p<0.001, ****p<0.0001.

Due to the premature death of many PBS-treated *Cln8*^*mnd*^ mice, only WT and scAAV9.pT-MecP2.CLN8-treated *Cln8*^*mnd*^ mice could be examined from 10-24 months of age. Over these time points, scAAV9.pT-MecP2.CLN8-treated *Cln8*^*mnd*^ mice had more ASM in the VPM/VPL and S1BF than their WT counterparts, but ASM levels never approached those observed in moribund (eight month-old) PBS-treated Cln8 *Cln8*^*mnd*^ mice, even at 24 months of age. These reductions in ASM are far greater than those documented for any other therapy in *Cln8*^*mnd*^ mice^17^.

Mitochondrial ATP synthase subunit C (SubC) is a major constituent of the storage material in NCLs^4^. SubC was elevated in the VPM/VPL and S1BF in PBS-treated Cln8 *Cln8*^*mnd*^ mice at two, four, six, and eight months of age, with large increases at the latest time point (Figure 3C and 3D). At all of these time points, VPM/VPL and S1BF SubC levels in WT and scAAV9.pT-MecP2.CLN8-treated *Cln8*^*mnd*^ mice were indistinguishable. As with ASM, only WT and scAAV9.pT-MecP2.CLN8-treated *Cln8*^*mnd*^ mice could be analyzed from 10-24 months of age. While scAAV9.pT-MecP2.CLN8-treated *Cln8*^*mnd*^ mice showed some elevation of SubC in the VPM/VPL and S1BF at these late time points, SubC levels never approached those observed in moribund PBS-treated *Cln8*^*mnd*^ mice. These drastic reductions we observed in cortical and thalamic ASM and SubC suggest cellular disease burden in the brain is greatly reduced in scAAV9.pT-MecP2.CLN8-treated *Cln8*^*mnd*^ mice.

### Gliosis is reduced in scAAV9.pT-MecP2.CLN8-treated *Cln8*^*mnd*^ mice

In neurodegenerative disorders including NCLs, glial cells respond to neuronal damage with a predictable cascade of morphological and molecular changes known as reactive gliosis^32^. In the *Cln8*^*mnd*^ mouse, brain gliosis has been reported to precede neuron loss, with microglial activation appearing prior to 3 months of age, followed shortly thereafter by astrocytosis^6^. We observed the first signs of reactive microglia, as evidenced by increased immunolabeling for cluster of differentiation 68 (CD68) in the VPM/VPL and S1BF of PBS-treated *Cln8*^*mnd*^ mice at two months of age (Figure 4A and 4B). In the subsequent months, CD68 immunoreactivity increased dramatically, with particularly large increases at the moribund time point of eight months of age. CD68-positive microglia in PBS-treated *Cln8*^*mnd*^ mice displayed morphology consistent with a reactive state, with hypertrophic soma and little ramification.

**Figure 4.**
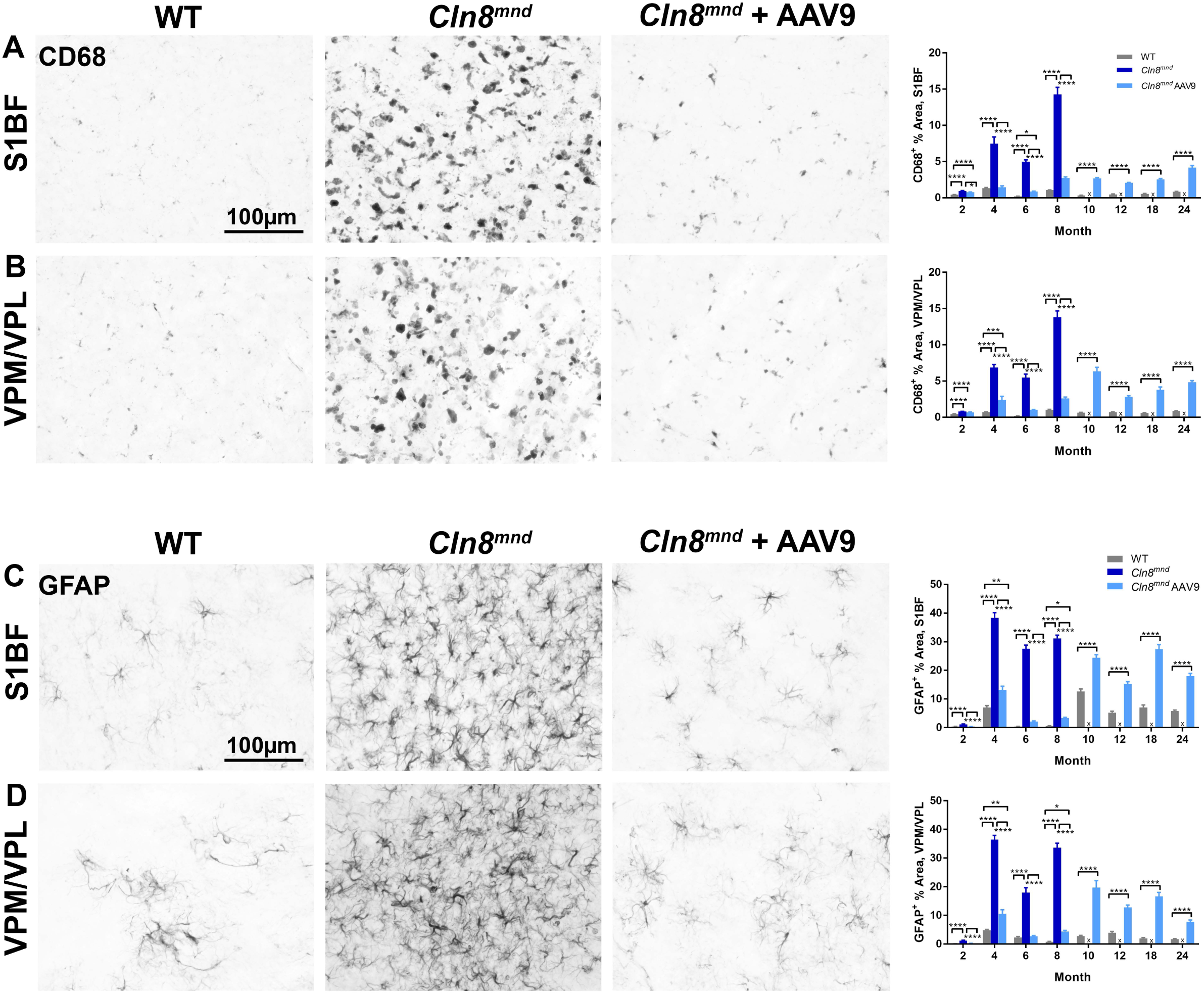
scAAV9.pT-MecP2.CLN8 treatment reduces glial activation in *Cln8*^*mnd*^ mice. As measured by immunohistochemistry, scAAV9.pT-MecP2.CLN8 treatment significantly prevents microglial activation (CD68; A,B) and astrocyte activation (GFAP; C,D) in the S1BF and VPM/VPL of 24 month old *Cln8*^*mnd*^ mice. Images representative of 8 months of age. Mean ± SEM. Each time point was analyzed independently as either a one-way ANOVA, Fisher’s LSD (2 to 8 months), or unpaired t-test (10 to 24 months). Biological N=4/sex/genotype. Detailed sample Ns are described in Supplemental Table 2. ***p<0.001, ****p<0.0001.

scAAV9.pT-MecP2.CLN8-treated *Cln8*^*mnd*^ mice had significantly less microglial activation than PBS-treated *Cln8*^*mnd*^ mice at all time points examined in the S1BF, and from four months of age onwards in the VPM/VPL. CD68 levels were indistinguishable from WT levels at several early (2-8 month) time points in scAAV9.pT-MecP2.CLN8-treated *Cln8*^*mnd*^ mice, although slight elevations were present at four and eight months of age. CD68 levels remained elevated in scAAV9.pT-MecP2.CLN8-treated *Cln8*^*mnd*^ mice through 24 months of age, but generally remained lower than the levels observed in moribund PBS-treated *Cln8*^*mnd*^ mice.

We observed the first signs of astrocytosis, as evidenced by slight increases in glial fibrillary acidic protein (GFAP) immunoreactivity, in PBS-treated *Cln8*^*mnd*^ mice at two months of age in the VPM/VPL and S1BF (Figure 4C and 4D). This was followed at four months of age by dramatic increases in GFAP immunoreactivity in both areas, indicating profound astrocytosis. Astrocytes in PBS-treated *Cln8*^*mnd*^ mice appeared hypertrophic and intensely GFAP-positive, typical of the reactive responses observed in neurodegenerative diseases. GFAP immunoreactivity remained highly elevated from four through eight months.

Treatment with scAAV9.pT-MecP2.CLN8 greatly reduced or eliminated astrocytosis, resulting in GFAP immunoreactivity indistinguishable from WT levels at two and six months of age, and far lower than PBS-treated *Cln8*^*mnd*^ mice at four and eight months of age, both in the VPM/VPL and S1BF. Moderate astrocytosis became evident from 10 through 24 months in the VPM/VPL and S1BF in scAAV9.pT-MecP2.CLN8-treated *Cln8*^*mnd*^ mice, but never exceeded that observed in moribund PBS-treated *Cln8*^*mnd*^ mice. Together, these results demonstrate that scAAV9.pT-MecP2.CLN8 administration greatly attenuates glial pathology in *Cln8*^*mnd*^ mice.

### scAAV9.pT-MecP2.CLN8 reduces motor and behavioral abnormalities in *Cln8*^*mnd*^ mice

Given the profound rescue we observed in lifespan and brain histopathological parameters in scAAV9.pT-MecP2.CLN8-treated *Cln8*^*mnd*^ mice, we anticipated improvements in behavioral measures of CNS function. We performed a battery of behavioral tests to comprehensively assess performance in motor and learning/memory-related tasks, focusing on behavioral parameters that have been shown to be disrupted in the *Cln8*^*mnd*^ model^18, 19, 24, 33^.

To quantify motor coordination, we measured Rotarod performance from 2-24 months of age. Rotarod performance declined rapidly in PBS-treated *Cln8*^*mnd*^ mice, with significant performance deficits evident at eight and ten months of age (Figure 5A). scAAV9.pT-MecP2.CLN8 administration completely rescued Rotarod performance at all time points, with scAAV9.pT-MecP2.CLN8-treated *Cln8*^*mnd*^ mice performing similarly to WT mice from 2-24 months of age. Similar motor coordination deficits were also evident in the vertical pole test, where PBS-treated *Cln8*^*mnd*^ mice exhibited increased descent time, time to turn down the pole, and fall frequency at eight and 10 months of age (Figure 5B-D). Treatment with scAAV9.pT-MecP2.CLN8 reduced these three deficits at both time points, with scAAV9.pT-MecP2.CLN8-treated animals maintaining WT levels of performance until 24 months of age.

**Figure 5.**
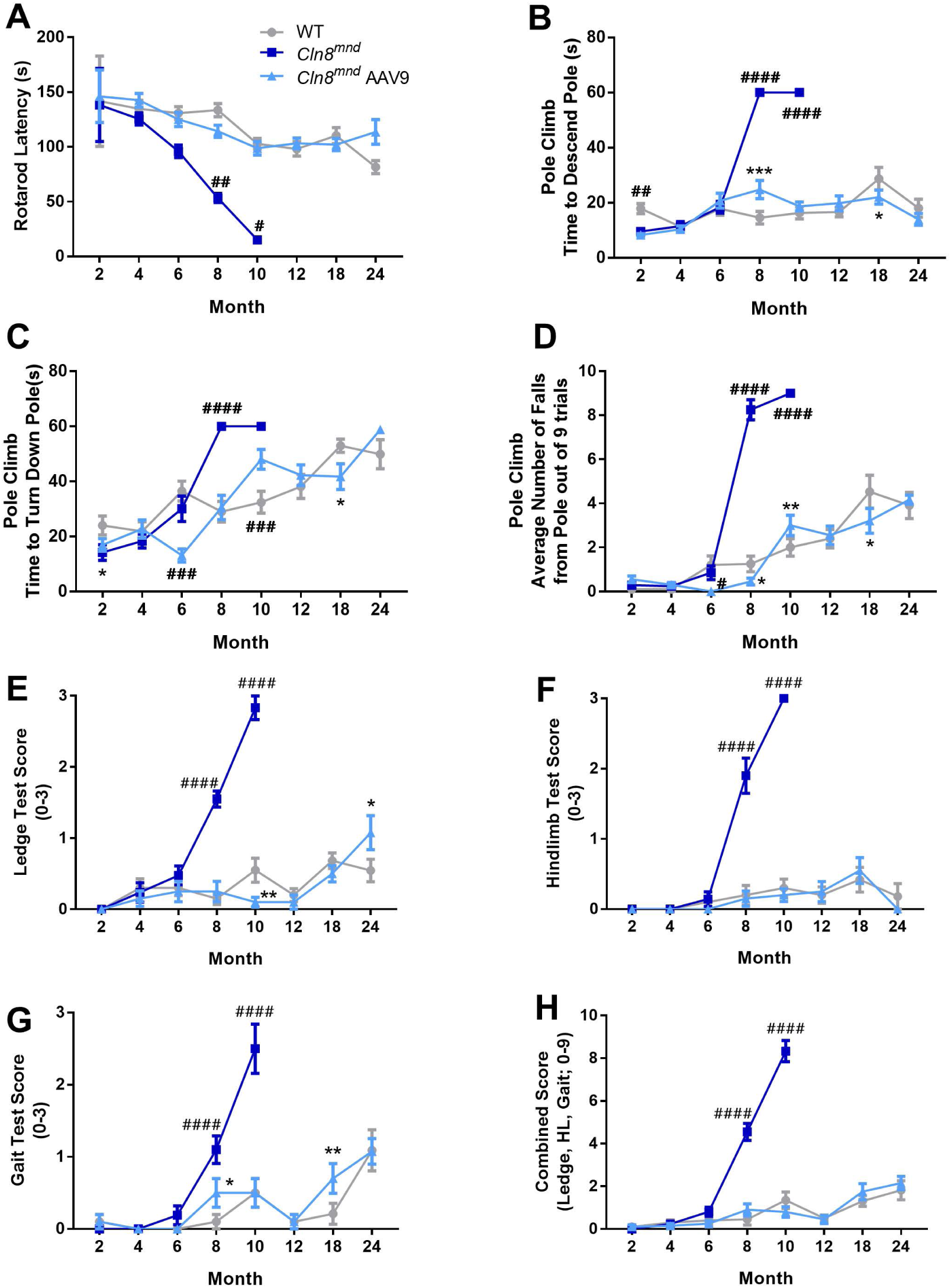
scAAV9.pT-MecP2.CLN8 prevents motor abnormalities in *Cln8*^*mnd*^ mice. Neonatal administration of scAAV9.pT-MecP2.CLN8 leads to prevention of motor abnormalities as measured by the accelerating Rotarod (A), vertical pole climb (B-D), and coordination tasks (E-H). Mean ± SEM, two-way ANOVA, Fisher’s LSD. Detailed Ns are described in Supplemental Table 2. Asterisks indicate difference from WT group: *p<0.05, **p<0.01, ***p<0.001. Hashes indicate differences from all other groups at that time point: #p<0.05, ##p<0.01, ###p<0.001, ####p<0.0001.

To further examine parameters of motor performance, we deployed a three component assay that has previously shown utility in mouse models of neurological disease^26, 34^. At eight and 10 months of age, PBS-treated *Cln8*^*mnd*^ mice showed impairments in hind limb clasping frequency, ledge descent ability, and overall gait, and had significantly worse combined scores (Figure 5E-H). scAAV9.pT-MecP2.CLN8-treated *Cln8*^*mnd*^ mice, however, performed at or near WT levels in all three tasks at both time points, with combined scores indistinguishable from WT at eight and 10 months of age. In all measures of motor performance (Rotarod, pole climb, and three component test), the behavioral improvements noted with scAAV9.pT-MecP2.CLN8 administration persisted well beyond the moribund time point of eight months for *Cln8*^*mnd*^ mice, with performance remaining at or near WT levels for scAAV9.pT-MecP2.CLN8-treated *Cln8*^*mnd*^ mice through 24 months of age.

To test whether learning and memory were similarly rescued with scAAV9.pT-MecP2.CLN8, we measured performance in the Morris Water Maze. During “training” trials, six and eight month-old PBS-treated *Cln8*^*mnd*^ mice took longer to swim to a platform marked by a visual cue, despite having similar swim speed (Figure 6A-B). scAAV9.pT-MecP2.CLN8 treatment rescued time to reach the platform to intermediate levels at both time points, though treated animals ultimately performed similarly to untreated animals as they aged. In “memory” trials in which the visual cue was removed from the platform, PBS-treated *Cln8*^*mnd*^ mice took longer to swim to the platform at six and eight months of age, but these results were confounded by differences in swim speed during these trials (Figure 6C-D). scAAV9.pT-MecP2.CLN8 treatment rescued eight month swim speed to WT levels, and six and eight month platform finding latency to intermediate levels. Finally, during intermittent “learning” (also known as “reversal”) trials, in which the unmarked platform was moved to a novel location, PBS-treated *Cln8*^*mnd*^ mice, which were only tested at 6 months of age as they perished by later trials, took longer to swim to the platform at 6 months of age, with no apparent rescue in the scAAV9.pT-MecP2.CLN8-treated group (Figure 6 E-F). Together, while the data suggest that some impairments in learning and memory may have been alleviated with scAAV9.pT-MecP2.CLN8 treatment, differences in motor performance and vision, as demonstrated by deficits in swim speed in the memory tests and deficits in time to the platform in the “training” trials, may have confounded the results.

**Figure 6.**
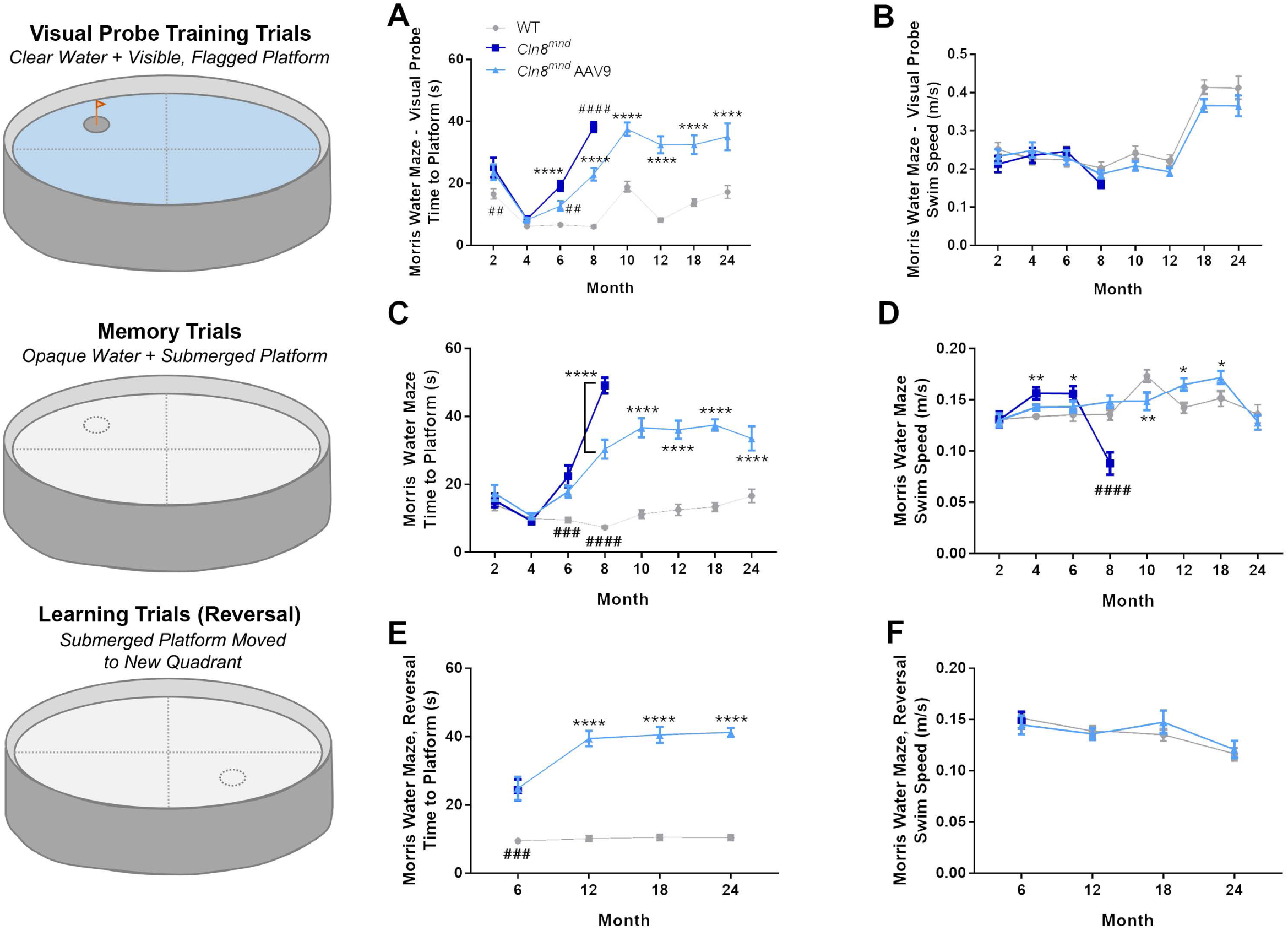
scAAV9.pT-MecP2.CLN8 does not prevent Morris water maze deficits in *Cln8*^*mnd*^ mice. Memory and learning abnormalities are partially, but not entirely, corrected as measured by the Morris water maze “training” (A-B), “memory” (C-D), and “learning” (“reversal”, E-F) tasks. Swim speeds are shown as a control, showing that motor deficits may play a role in water maze deficits. Mean ± SEM, two-way ANOVA, Fisher’s LSD. Detailed Ns are described in Supplemental Table 2. Asterisks indicate difference from WT group: *p<0.05, **p<0.01, ****p<0.0001. Hashes indicate differences from all other groups at that time point: ##p<0.01, ###p<0.001, ####p<0.0001.

While caring for these mice, we were surprised to observe tremors in many moribund PBS-treated *Cln8*^*mnd*^ mice, a phenotype that mirrors symptoms observed in some human CLN8 disease patients^35^, but has not been reported in the *Cln8*^*mnd*^ literature. To quantify the phenotype, we measured tremor scores in a force plate actimeter. Force plate data showed that scAAV9.pT-MecP2.CLN8-treated mice maintained a healthy weight through 24 months of age, and showed no consistent differences in force plate parameters of total distance travelled, bouts of low mobility, or total area covered, but exhibited slightly less focused stereotypy as they aged. (Figure 7A-F). PBS-treated *Cln8*^*mnd*^ mice exhibited progressive elevations in tremor scores beginning at 2 months of age, which culminated in large elevations in tremor scores at ten months of age (Figure 7G-J). Treatment with scAAV9.pT-MecP2.CLN8 prevented the development of increased tremor scores at most time points and frequencies. Notably, even at the last time point examined (24 months), scAAV9.pT-MecP2.CLN8-treated *Cln8*^*mnd*^ mice failed to exhibit the large elevations in tremor scores evident in moribund PBS-treated *Cln8*^*mnd*^ mice. Together, these results demonstrate unprecedented improvements in survival, brain pathology, and behavioral dysfunction in *Cln8*^*mnd*^ mice following a single postnatal ICV injection of scAAV9.pT-MecP2.CLN8.

**Figure 7.**
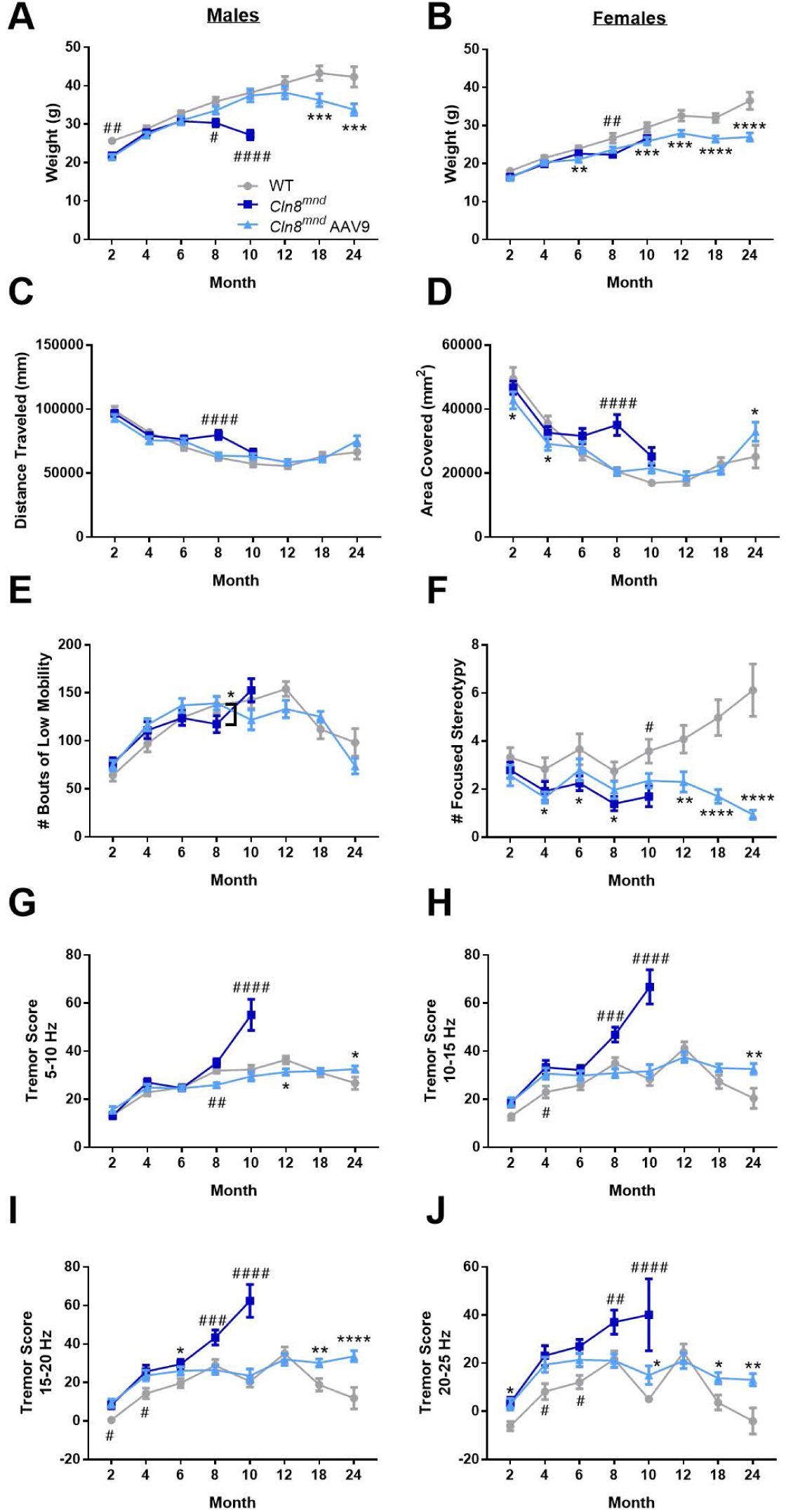
scAAV9.pT-MecP2.CLN8 prevents tremor presentations in *Cln8*^*mnd*^ mice. scAAV9.pT-MecP2.CLN8 treated mice show slight differences in weight, but maintain a healthy weight through 24 months of age (A, males; B, females), and show no consistent differences in force plate parameters of total distance travelled (C), total area covered (D), and bouts of low mobility (E). scAAV9.pT-MecP2.CLN8 treated mice show slight, but significantly fewer stereotypic movements as they age (F). Lastly, AAV9 treatment prevents tremor presentation in *Cln8*^*mnd*^ mice at various frequencies through 24 months of age, though AAV9 treated mice show slightly more tremors than their wildtype counterparts as they age (G-J). Mean ± SEM, two-way ANOVA, Fisher’s LSD. Detailed Ns are described in Supplemental Table 2. Asterisks indicate difference from WT group: *p<0.05, **p<0.01, ***p<0.001, ****p<0.0001. Hashes indicate differences from all other groups at that time point: #p<0.05, ##p<0.01, ###p<0.001, ####p<0.0001.

## Discussion

CLN8 disease is a phenotypically variable, but often severe form of NCL with no known cure or treatment. While some preclinical studies have shown positive effects following the administration of small molecule therapies, behavioral and pathological improvements have been modest^18, 19^, and lifespan extension has been minimal^17^. To date, the paucity of effective treatments is reflected in the clinic, with no clinical trials registered at clinicaltrials.gov to test any therapy in CLN8 disease patients.

As is the case with many other NCLs, the lack of therapies can be attributed to a poor understanding of CLN8 function. Recent studies have made great progress towards this end, demonstrating that CLN8 binds directly to both soluble lysosomal enzymes and COPI/COPII machinery, facilitating the ER to Golgi trafficking of lysosomal enzymes^5^. Still, while this knowledge provides valuable insight into potential drug targets, it is only a first step in an often long and arduous pathway towards the development of a small molecule therapy. Treatments that can directly resolve CLN8 deficiency are desperately needed.

Recent clinical trials for NCLs have focused on two primary treatment modalities: enzyme replacement therapy (ERT) and gene therapy, both of which aim to restore functional CLN proteins in the most affected cell populations (i.e. neurons). ERT has been used to deliver recombinant proteins directly into the brain through ICV injection. This approach has been very successful in some cases; cerliponase alfa (trade name Brineura) was FDA approved in 2017 for the treatment of CLN2 disease, which is caused by a deficiency in the soluble lysosomal enzyme tripeptidyl peptidase-1 (TPP1). Cerliponase alfa is a recombinant proenzyme form of endogenous TPP1, and while the therapy slows disease progression, it must be administered biweekly for a patient’s entire remaining lifespan, and adverse events such as hypersensitivity reactions are common^36^. Enzyme replacement therapy is even less appealing for some other NCLs, particularly those caused by deficiencies in larger transmembrane proteins that would be more difficult to target to the correct subcellular locations.

Gene therapy offers another method to restore functional NCL proteins in vulnerable cell populations. With these therapies, viral vectors are used to drive expression of the gene of interest, with AAVs being the vector of choice in recent preclinical NCL studies^26, 37-41^. Different AAV serotypes have varying tropism and distribution patterns, allowing for specific tissues and cell populations to be targeted. AAV9 is particularly useful for CNS disorders, as it crosses the blood brain barrier following intravenous administration to target both neurons and glia with high efficiency^27-29, 42^. Despite this useful property, other more direct routes of administration to the CNS are often still preferred since doses can be decreased, minimizing the potential for hepatotoxicity and other peripheral side effects^43^. ICV delivery facilitates much smaller doses while still transducing CNS neurons with high efficiency, and while our study utilized ICV delivery for practicality in mice, the results are supportive of other CSF-mediated delivery methods that are currently used in patients, such as intrathecal delivery^26^.

Here, we demonstrate that a single P1 injection of an AAV9 viral vector driving expression of human *CLN8* rescues many of the pathological and behavioral hallmarks of the disease in a mouse model of CLN8 disease. Our vector, scAAV9.pT-MecP2.CLN8, utilizes a truncated form of the *MecP2* promoter in order to achieve physiologically appropriate expression levels^29^ in transduced cells. We administered the virus at P1, since this timing in mice best recapitulates the transduction patterns observed in non-human primates and anticipated in human patients^44-46^. This dosing scheme would be most representative of early postnatal treatment in humans rather than symptomatic stages, and future adoption of newborn screening for *CLN8* mutations could make this type of treatment both realistic and feasible.

This single P1 ICV injection of scAAV9.pT-MecP2.CLN8 achieved widespread and persistent expression of *hCLN8* in the CNS from 4-24 months of age. Restoration of functional CLN8 greatly reduced histopathological hallmarks of the disease, including ASM, SubC accumulation, microglial activation, and astrocytosis, providing protection through 24 months of age (the final time point assessed). The therapy also rescued a number of behavioral parameters, including motor performance in Rotarod, pole climb, and three component tests. Morris Water Maze test performance was improved in scAAV9.pT-MecP2.CLN8-treated *Cln8*^*mnd*^ mice, although this test may have been influenced by differences in motor performance. Additionally, while we did not measure retinal function or visual acuity, deficits have been documented for *Cln8*^*mnd*^ mice at as young as two months of age. Thus, it is possible that differences (and/or corrections) in visual function accounted for some of the behavioral results we observed.

We also documented, for the first time, the presence of tremors in *Cln8*^*mnd*^ mice. scAAV9.pT-MecP2.CLN8 treatment reduced tremor scores from two through 18 months, with scores in scAAV9.pT-MecP2.CLN8-treated *Cln8*^*mnd*^ mice indistinguishable from WT at most frequencies and time points. Finally, and perhaps the most encouraging result in our study, was the increased survival conferred to *Cln8*^*mnd*^ mice by scAAV9.pT-MecP2.CLN8 treatment. While PBS-treated *Cln8*^*mnd*^ mice all died by 12 months of age, scAAV9.pT-MecP2.CLN8-treated animals survived for the duration of the study (24 months), with survival curves indistinguishable from WT mice.

While these results are encouraging, several important questions remain. At later time points beyond eight months, increased lysosomal storage material and gliosis were evident in the brains of scAAV9.pT-MecP2.CLN8-treated *Cln8*^*mnd*^ mice. While levels never approached those observed in PBS-treated *Cln8*^*mnd*^ mice, this suggests that the rescue obtained by a single P1 ICV injection of scAAV9.pT-MecP2.CLN8 is incomplete. Whether these minor tissue-level defects reflect incomplete rescue in transduced cells or are the consequences of cellular dysfunction in untransduced cells such as microglia is currently unknown. Further experiments will be necessary to answer these questions and to further enhance the efficacy of the therapy. Collectively, the improvements in survival, pathology, motor function, and behavior obtained from scAAV9.pT-MecP2.CLN8 administration in this study far surpassed those reported for any other therapy in *Cln8*^*mnd*^ mice, suggesting great potential for a similar strategy in human CLN8 patients.

## Acknowledgments/Funding

This work received support from the Cure Batten CLN8 Velona Foundation, Amicus Therapeutics, Sanford Research Imaging Core within the Sanford Research Center for Pediatric Research (NIH P20GM103620), and the Sanford Research Molecular Pathology Core within the Sanford Research Center for Cancer Biology (NIH P20GM103548).

## Author Contributions

Conceptualization: TBJ, JTC, KM, JMW; Methodology: TBJ, JTC, MAP, CDB; Formal Analysis: TBJ, KAW, JTC, LL, MAP; Investigation: TBJ, LL, MAP, CDB, DJT, SSD, BM, SL; Resources: KM; Writing – Original Draft: JB; Writing – Review and Editing: TBJ, KAW, JTC, JB, KM, JMW; Visualization: TBJ, KAW; Supervision: TBJ, JTC, KM, JMW; Project Administration: TBJ, KAW, JMW; Funding Acquisition: JMW.

## Materials & Methods

### Ethics Statement/Animals

Wild type and homozygous *Cln8*-mutant mice (*Cln8*^*mnd*^) on C57BL/6J backgrounds were used for all studies. *Cln8*^*mnd*^ mice harbor a single nucleotide insertion (267-268C, codon 90), predicting a frameshift and premature STOP codon. All mice were housed under identical conditions in an AAALAC accredited facility with IACUC approval (USDA License 46-R-0009; Protocol #156-01-22D, Sanford Research, Sioux Falls, SD).

### Vector and AAV9 Delivery

A human CLN8 cDNA clone was subcloned into an AAV vector under the control of a truncated *MecP2* (pT-MecP2) promoter. Self-complementary AAV9.pT-MecP2.CLN8 was produced as previously described, using transient transfection procedures with a double-stranded AAV2-ITR–based pT-MecP2.CLN8 vector, with a plasmid encoding Rep2Cap9 sequence along with an adenoviral helper plasmid pHelper (Stratagene, Santa Clara, CA) in HEK293 cells^42^. Vectors were purified by double cesium chloride gradient centrifugation. Silver staining analysis was used to measure the purity and titer of the vector.

At P0 or P1, mouse pups were sexed then randomly assigned to a treatment group of either PBS- or AAV9.pT-MecP2.CLN8-treated. Pups were randomly assigned until all treatment groups were filled. Mice received a single intracerebroventricular (ICV) injection of either PBS or scAAV9.pT-MecP2.CLN8 at a dose of 5.0 x 10^10^ vg/animal. Animals were anesthetized with hypothermia at P1, injected with 4 μL of solution, and monitored until fully recovered. All animals were monitored daily by trained animal technicians and genotyped as previously described^26^.

### qPCR

Mice were CO_2_ euthanized and a 3mm lateral section of the outer right hemisphere was collected and frozen for RNA isolation. Total RNA was extracted according to the manufacturer’s protocol for the Maxwell 16 LEV simplyRNA Tissue Kit (AS1280), using a Maxwell 16 MDx machine (Promega). RNA quality and concentration were assayed using a BioTek Epoch Microplate Spectrophotometer, with all samples having concentrations between 200-1000 ng/μl and A260/A280 > 2. Using the Promega GoScript Reverse Transcription System (A5001) and the manufacturer’s protocol, cDNA synthesis was performed on 1μg of total RNA. The following primers were used from Integrated DNA Technologies: *hCLN8* Forward: GGACTGGCTCTGCTTACGCTAA; Reverse: GCTCTTGGCTTCTGGCTGTG. *Gapdh* Forward: ACC ACA GTC CAT GCC ATC AC; Reverse: TCC ACC ACC CTG TTG CTG TA. Reaction were run on an Applied Biosystems ABI 7500 Fast Real-time PCR System, and PCR products were run on 3% TBE gels at 125 V for 90 minutes. Bands were visualized under ultraviolet light using a Gel-Doc Imager (UVP), and densitometry was performed with VisionWorks software (UVP). Relative levels of scAAV9.pT-MecP2. CLN8 transcript levels were normalized against *Gapdh*.

### RNAscope

hCln8 transcript localization was visualized with RNAscope as previously described^26^. Briefly, brains were collected from CO_2_ euthanized mice and placed on a 1mm sagittal brain block. Tissue blocks from 0 to 3mm right of the midline were flash frozen. Brain sections were sliced on a cryostat at 16 μm, series dehydrated, placed on slides, and processed for RNAscope according to the manufacturer’s protocols. Sections were labeled with a human *CLN8* probe (ACDBio Cat No 516851), counterstained with DAPI, and mounted on slides using antifade mounting media (Dako faramount, Agilent).

### Immunohistochemistry

The remaining left hemisphere of the brain was fixed in 4% paraformaldehyde and sectioned into 50 μm slices with selected slices placed into a 24 well plate. Immunohistochemistry was performed on free-floating sections as previously described using anti-ATP synthase subunit C (Abcam, ab181243), anti-GFAP (Dako, Z0334), and anti-CD68 (AbD Serotec, MCA1957) antibodies^26^. Secondary antibodies included anti-rat and anti-rabbit biotinylated (Vector Labs, BA-9400). Sections were placed on slides and mounted using xylene and xylene-based mounting media (DPX, VWR International). Sections were imaged and analyzed using an Aperio Digital Pathology Slide Scanner (VERSA) and associated software. Multiple images of each animal were taken in the VPM/VPL of the thalamus and the S1BF, and images were quantified using ImageJ.

### Neurobehavior testing

#### Rotarod

Animals participated in an accelerated Rotarod protocol as previously described to assess motor coordination (Columbus Instruments, Columbus, OH, USA)^26^. The machine was set to accelerate 0.3 rpm every two seconds, with a starting speed of 0.3 rpm and a maximum speed of 36 rpm. Mice were trained for three consecutive trials, given a 30 min rest period, trained for three consecutive trials, given a second 30 min rest period, and trained for three final consecutive trials. After a four-hour rest period, mice were tested using the same paradigm as the training session. The latency time to fall from the rod was averaged from each of the nine afternoon testing sessions to produce one value per mouse.

#### Pole Climb

The pole climb descent test was performed as previously described^47^. Mice were placed downward on a metal pole and given 60 seconds to descend the pole. Mice were placed upward on a metal pole and given 60 seconds to turn downward on the pole. Lastly, the number of falls made by each mouse during the two tests was recorded.

#### Water Maze

Mice were tested in a 4 foot diameter Morris Water Maze apparatus to assess memory and learning deficiencies as previously described^26^. The apparatus was filled with water to ∼26 inches and the goal platform submerged by 0.5cm at 315°. The tub was aligned with four distinct visual cues at 0, 90, 180, and 270° to aid in spatial memory. Mice were first trained in a clear pool with a flagged platform to rule out confounding factors such as visual, motor, and anxious difficulties. Mice were given 60 seconds to find the platform each trial, with four trials in the morning, followed by a three-hour rest period, and four additional trials in the afternoon. Mice that could not locate the platform with 50% accuracy in the time allotted were eliminated from further testing. Mice were then tested in water colored with white, non-toxic tempura paint and an un-flagged platform. Mice were given 60 seconds to complete each trial, with four trials in the morning, followed by a three-hour rest period, and four additional trials in the afternoon. Mice were tested for four consecutive days, each day starting at a different visual cue. Mice were recorded using Any-maze video tracking software (Stoelting Co., Wood Dale, IL, USA). Test duration and swim speed were represented as the average from the sixteen afternoon trials per mouse. At 6, 12, 18, and 24 months of age, an additional four days of reversal testing were introduced, with the hidden platform moved from 315 degrees to 45 degrees.

#### Clasping, Ledge, and Gait Tests

Tests were performed as previously described^26, 34^. For hind limb clasping measurements, animals were scored on the extent to which their limbs clasped into their abdomen when held by the base of their tail (score 0-3). For ledge lowering measurements, animals were scored on their ability to climb down from the edge of their home cage (score 0-3). For gait measurements, animals were scored on their overall ease of walking, including whether their abdomen dragged on the ground and if their limbs were splayed out while walking (score 0-3). The same blinded experimenter determined all scores.

#### Force Plate

A force plate actimeter was used to measure tremors as previously described^47^. Animals were recorded in a sound-proof chamber for 20 minutes, and data was processed using FPA Analysis Software (BASi, West Lafayette, IN).

### Statistical Analysis

Statistical analyses were performed using GraphPad Prism (v6.04+) and details are noted in the figure legends. In general, one-way ANOVA was employed with Fisher’s LSD, and outliers were removed with the ROUT method, Q=0.1%. If appropriate, an unpaired t-test was used. For the survival curve analysis, the log-rank (Mantel-Cox) test was used. *p<0.05, **p<0.01, ***p<0.001, ****p<0.0001.

