## Supplemental Tables 1-2 for "AAV9 gene therapy restores lifespan and treats pathological and behavioral abnormalities in a mouse model of CLN8-Batten disease"

**Supplemental Table 1. Death and euthanasia descriptions.**

| Treatment Group | Sex | Age at Death | Morbidity Notes |
|---|---|---|---|
| WT | F | 16 | Found dead - likely intestinal tumor leading to rupture of blood vessel and internal hemorrhaging |
| WT | F | 17 | Euthanized - significantly underweight, labored breathing, lateral recumbancy, hunched |
| WT | F | 22 | Euthanized - labored breathing; moribund |
| WT | F | 24 | n/a - retired from study |
| WT | F | 24 | n/a - retired from study |
| WT | F | 24 | n/a - retired from study |
| WT | F | 24 | n/a - retired from study |
| WT | F | 24 | n/a - retired from study |
| WT | F | 24 | n/a - retired from study |
| WT | F | 24 | n/a - retired from study |
| WT | F | 24 | n/a - retired from study |
| WT | F | 24 | n/a - retired from study |
| WT | F | 24 | n/a - retired from study |
| WT | F | 24 | n/a - retired from study |
| WT | F | 24 | n/a - retired from study |
| WT | M | 24 | n/a - retired from study |
| WT | M | 24 | n/a - retired from study |
| WT | M | 24 | n/a - retired from study |
| WT | M | 24 | n/a - retired from study |
| WT | M | 24 | n/a - retired from study |
| WT | M | 24 | n/a - retired from study |
| WT | M | 24 | n/a - retired from study |
| WT | M | 24 | n/a - retired from study |
| WT | M | 24 | n/a - retired from study |
| WT | M | 24 | n/a - retired from study |
| WT | M | 24 | n/a - retired from study |
| WT | M | 24 | n/a - retired from study |
| WT | M | 24 | n/a - retired from study |
| WT | M | 24 | n/a - retired from study |
| WT | M | 24 | n/a - retired from study |
| $Cln8^{mnd}$ + AAV9 | F | 17 | Euthanized - prolapsed/bleeding rectum |
| $Cln8^{mnd}$ + AAV9 | F | 18 | Euthanized - Significant growth appearing on lateral abdomen, emaciation, impaired movement due to growth |
| $Cln8^{mnd}$ + AAV9 | F | 24 | n/a - retired from study |

| | | | |
|---|---|---|---|
| *Cln8*$^{mnd}$ + AAV9 | F | 24 | n/a - retired from study |
| *Cln8*$^{mnd}$ + AAV9 | F | 24 | n/a - retired from study |
| *Cln8*$^{mnd}$ + AAV9 | F | 24 | n/a - retired from study |
| *Cln8*$^{mnd}$ + AAV9 | F | 24 | n/a - retired from study |
| *Cln8*$^{mnd}$ + AAV9 | F | 24 | n/a - retired from study |
| *Cln8*$^{mnd}$ + AAV9 | F | 24 | n/a - retired from study |
| *Cln8*$^{mnd}$ + AAV9 | F | 24 | n/a - retired from study |
| *Cln8*$^{mnd}$ + AAV9 | F | 24 | n/a - retired from study |
| *Cln8*$^{mnd}$ + AAV9 | F | 24 | n/a - retired from study |
| *Cln8*$^{mnd}$ + AAV9 | F | 24 | n/a - retired from study |
| *Cln8*$^{mnd}$ + AAV9 | F | 24 | n/a - retired from study |
| *Cln8*$^{mnd}$ + AAV9 | F | 24 | n/a - retired from study |
| *Cln8*$^{mnd}$ + AAV9 | M | 23 | Euthanized - abdominal mass; mobility issues |
| *Cln8*$^{mnd}$ + AAV9 | M | 24 | n/a - retired from study |
| *Cln8*$^{mnd}$ + AAV9 | M | 24 | n/a - retired from study |
| *Cln8*$^{mnd}$ + AAV9 | M | 24 | n/a - retired from study |
| *Cln8*$^{mnd}$ + AAV9 | M | 24 | n/a - retired from study |
| *Cln8*$^{mnd}$ + AAV9 | M | 24 | n/a - retired from study |
| *Cln8*$^{mnd}$ + AAV9 | M | 24 | n/a - retired from study |
| *Cln8*$^{mnd}$ + AAV9 | M | 24 | n/a - retired from study |
| *Cln8*$^{mnd}$ + AAV9 | M | 24 | n/a - retired from study |
| *Cln8*$^{mnd}$ + AAV9 | M | 24 | n/a - retired from study |
| *Cln8*$^{mnd}$ + AAV9 | M | 24 | n/a - retired from study |
| *Cln8*$^{mnd}$ + AAV9 | M | 24 | n/a - retired from study |
| *Cln8*$^{mnd}$ + AAV9 | M | 24 | n/a - retired from study |
| *Cln8*$^{mnd}$ + AAV9 | M | 24 | n/a - retired from study |
| *Cln8*$^{mnd}$ + AAV9 | M | 24 | n/a - retired from study |
| *Cln8*$^{mnd}$ + AAV9 | M | 24 | n/a - retired from study |
| *Cln8*$^{mnd}$ | F | 7 | Euthanized - severe ulcerative dermatitis |
| *Cln8*$^{mnd}$ | F | 9 | Euthanized - seizures, severely impaired ambulation |
| *Cln8*$^{mnd}$ | F | 9 | Euthanized - seizures, severely impaired ambulation |
| *Cln8*$^{mnd}$ | F | 9 | Found dead |
| *Cln8*$^{mnd}$ | F | 9 | Euthanized - extreme weight loss |
| *Cln8*$^{mnd}$ | F | 9 | Euthanized - humane end point |
| *Cln8*$^{mnd}$ | F | 9 | Found dead |

| | | | |
|---|---|---|---|
| *Cln8*$^{mnd}$ | F | 10 | Found dead |
| *Cln8*$^{mnd}$ | F | 10 | Euthanized - did not recover from severe seizures |
| *Cln8*$^{mnd}$ | F | 10 | Found dead |
| *Cln8*$^{mnd}$ | F | 10 | Euthanized - ataxia, inability to right from back, weight loss, little control of back legs |
| *Cln8*$^{mnd}$ | F | 10 | Found dead |
| *Cln8*$^{mnd}$ | F | 10 | Euthanized - humane end point |
| *Cln8*$^{mnd}$ | F | 11 | Euthanized - ataxia, significant weight loss, extreme diffculty in movement |
| *Cln8*$^{mnd}$ | M | 9 | Found dead |
| *Cln8*$^{mnd}$ | M | 9 | Found dead |
| *Cln8*$^{mnd}$ | M | 9 | Euthanized - seizures, severely impaired ambulation |
| *Cln8*$^{mnd}$ | M | 9 | Found dead |
| *Cln8*$^{mnd}$ | M | 10 | Euthanized - humane end point |
| *Cln8*$^{mnd}$ | M | 10 | Found dead |
| *Cln8*$^{mnd}$ | M | 10 | Found dead |
| *Cln8*$^{mnd}$ | M | 10 | Euthanized - humane end point |
| *Cln8*$^{mnd}$ | M | 10 | Euthanized - humane end point |
| *Cln8*$^{mnd}$ | M | 10 | Euthanized - humane end point |
| *Cln8*$^{mnd}$ | M | 10 | Euthanized - humane end point |
| *Cln8*$^{mnd}$ | M | 10 | Euthanized - ataxia, difficulty getting food and water, dropping weight, rough coat |
| *Cln8*$^{mnd}$ | M | 11 | Euthanized - unable to right itself, twitchy and hunched appearance |
| *Cln8*$^{mnd}$ | M | 12 | Euthanized - humane end point |
| *Cln8*$^{mnd}$ | M | 12 | Euthanized - hunched rough coat, priapism, general dyskinesia |
| *Cln8*$^{mnd}$ | M | 12 | Euthanized - hunched rough coat, priapism, general dyskinesia |

**Supplemental Table 2. Detailed Ns.**

| | | WT | | | | | | | | *Cln8*$^{mnd}$ | | | | | | | | *Cln8*$^{mnd}$ + AAV9 | | | | | | | |
|---|---|---|---|---|---|---|---|---|---|---|---|---|---|---|---|---|---|---|---|---|---|---|---|---|---|
| | | 2 | 4 | 6 | 8 | 10 | 12 | 18 | 24 | 2 | 4 | 6 | 8 | 10 | 12 | 18 | 24 | 2 | 4 | 6 | 8 | 10 | 12 | 18 | 24 |
| qPCR | Cerebral Cortex | 4 | 5 | 6 | 5 | 6 | 5 | 4 | 6 | 5 | 6 | 4 | 5 | - | - | - | - | 7 | 6 | 5 | 6 | 7 | 6 | 5 | 5 |
| | Cervical SC | 3 | 6 | 6 | 6 | 6 | 5 | 5 | 5 | 5 | 6 | 5 | 6 | - | - | - | - | 5 | 6 | 5 | 5 | 4 | 6 | 6 | 6 |
| | Thoracic SC | 1 | 6 | 6 | 6 | 6 | 6 | 4 | 6 | 1 | 5 | 5 | 6 | - | - | - | - | 1 | 5 | 5 | 5 | 7 | 5 | 6 | 6 |
| | Lumbar SC | 3 | 6 | 6 | 4 | 6 | 6 | 3 | 5 | 2 | 6 | 5 | 3 | - | - | - | - | 2 | 6 | 5 | 4 | 6 | 6 | 5 | 6 |
| | Kidney | 4 | 6 | 5 | 3 | 6 | 4 | 5 | 4 | 4 | 5 | 5 | 4 | - | - | - | - | 4 | 6 | 4 | 6 | 7 | 5 | 7 | 6 |
| ASM | S1BF | 8 | 20 | 44 | 41 | 33 | 24 | 28 | 44 | 20 | 22 | 48 | 44 | - | - | - | - | 21 | 21 | 47 | 45 | 53 | 20 | 32 | 40 |
| | VPM/VPL | 8 | 19 | 47 | 43 | 33 | 24 | 32 | 48 | 20 | 24 | 47 | 48 | - | - | - | - | 24 | 18 | 48 | 41 | 55 | 20 | 32 | 40 |
| SubC | S1BF | 33 | 54 | 68 | 57 | 37 | 43 | 42 | 52 | 24 | 41 | 68 | 72 | - | - | - | - | 35 | 33 | 69 | 64 | 50 | 66 | 68 | 72 |
| | VPM/VPL | 35 | 52 | 69 | 56 | 42 | 38 | 38 | 61 | 15 | 44 | 68 | 72 | - | - | - | - | 26 | 32 | 62 | 66 | 56 | 67 | 68 | 62 |
| GFAP | S1BF | 48 | 50 | 63 | 62 | 60 | 60 | 45 | 68 | 38 | 51 | 72 | 72 | - | - | - | - | 40 | 48 | 69 | 68 | 36 | 72 | 68 | 72 |
| | VPM/VPL | 46 | 50 | 68 | 63 | 58 | 58 | 46 | 62 | 39 | 48 | 64 | 72 | - | - | - | - | 35 | 48 | 69 | 68 | 36 | 66 | 72 | 68 |
| CD68 | S1BF | 54 | 53 | 69 | 72 | 46 | 57 | 48 | 63 | 51 | 34 | 72 | 72 | - | - | - | - | 45 | 24 | 68 | 68 | 56 | 68 | 62 | 65 |
| | VPM/VPL | 54 | 54 | 70 | 72 | 48 | 57 | 48 | 68 | 45 | 33 | 63 | 72 | - | - | - | - | 42 | 27 | 72 | 68 | 60 | 67 | 68 | 60 |
| | Rotarod | 20 | 20 | 19 | 20 | 20 | 20 | 19 | 11 | 21 | 21 | 20 | 20 | 5 | - | - | - | 20 | 20 | 20 | 20 | 20 | 20 | 20 | 11 |
| Pole Climb | Down | 20 | 19 | 20 | 19 | 20 | 20 | 19 | 11 | 18 | 18 | 19 | 17 | 9 | - | - | - | 18 | 18 | 20 | 20 | 20 | 20 | 19 | 12 |
| | Turn | 20 | 20 | 20 | 20 | 20 | 20 | 18 | 11 | 20 | 20 | 20 | 19 | 8 | - | - | - | 20 | 20 | 18 | 18 | 20 | 20 | 19 | 12 |
| | Falls | 20 | 20 | 20 | 20 | 20 | 20 | 19 | 11 | 21 | 21 | 20 | 20 | 8 | - | - | - | 20 | 20 | 19 | 20 | 20 | 20 | 19 | 12 |
| Water Maze | Time | 19 | 19 | 20 | 19 | 19 | 20 | 19 | 10 | 15 | 15 | 16 | 10 | - | - | - | - | 15 | 15 | 15 | 15 | 12 | 10 | 9 | 6 |
| | Speed | 20 | 20 | 20 | 20 | 20 | 20 | 19 | 10 | 17 | 61 | 16 | 10 | - | - | - | - | 15 | 14 | 15 | 14 | 12 | 10 | 9 | 6 |
| | Reversal Time | - | - | 18 | - | - | 20 | 19 | 9 | - | - | 16 | - | - | - | - | - | - | - | 15 | - | - | 10 | 9 | 6 |
| | Reversal Speed | - | - | 20 | - | - | 20 | 18 | 10 | - | - | 16 | - | - | - | - | - | - | - | 15 | - | - | 10 | 9 | 6 |
| Coordination | Clasping | 20 | 20 | 20 | 20 | 20 | 20 | 19 | 11 | 21 | 21 | 21 | 20 | 6 | - | - | - | 20 | 20 | 20 | 20 | 20 | 20 | 20 | 13 |
| | Ledge | 20 | 20 | 20 | 20 | 20 | 20 | 19 | 11 | 21 | 21 | 21 | 20 | 6 | - | - | - | 20 | 20 | 20 | 20 | 20 | 20 | 20 | 13 |
| | Gait | 20 | 20 | 20 | 20 | 20 | 20 | 19 | 11 | 21 | 21 | 21 | 20 | 6 | - | - | - | 20 | 20 | 20 | 20 | 20 | 20 | 20 | 13 |
| | Combined Score | 20 | 20 | 20 | 20 | 20 | 20 | 19 | 11 | 21 | 21 | 21 | 20 | 6 | - | - | - | 20 | 20 | 20 | 20 | 20 | 20 | 20 | 13 |
| Force Plate | Male Weight | 8 | 10 | 10 | 10 | 10 | 10 | 10 | 7 | 10 | 10 | 10 | 10 | 5 | - | - | - | 11 | 10 | 10 | 10 | 10 | 10 | 10 | 6 |
| | Female Weight | 10 | 10 | 10 | 10 | 10 | 10 | 9 | 4 | 10 | 11 | 11 | 10 | 1 | - | - | - | 9 | 9 | 10 | 10 | 10 | 10 | 10 | 6 |
| | Total Distance | 19 | 20 | 20 | 20 | 20 | 20 | 19 | 11 | 20 | 21 | 21 | 20 | 6 | - | - | - | 20 | 20 | 20 | 19 | 19 | 19 | 20 | 12 |
| | Bouts Low Mobility | 19 | 20 | 20 | 20 | 20 | 20 | 19 | 11 | 20 | 21 | 21 | 20 | 6 | - | - | - | 20 | 20 | 20 | 20 | 20 | 20 | 20 | 12 |
| | Total Area | 19 | 20 | 20 | 20 | 19 | 20 | 19 | 11 | 20 | 21 | 21 | 19 | 6 | - | - | - | 20 | 20 | 20 | 18 | 19 | 19 | 19 | 12 |
| | Focused Stereo | 19 | 20 | 20 | 19 | 20 | 20 | 18 | 11 | 20 | 21 | 20 | 20 | 6 | - | - | - | 20 | 19 | 19 | 18 | 20 | 18 | 20 | 12 |

| | | | | | | | | | | | | | | | | | | | | | | | | | |
|---|---|---|---|---|---|---|---|---|---|---|---|---|---|---|---|---|---|---|---|---|---|---|---|---|---|
| | Tremor 5-10Hz | 20 | 20 | 20 | 20 | 20 | 20 | 19 | 11 | 21 | 21 | 21 | 20 | 6 | - | - | - | 20 | 20 | 20 | 20 | 20 | 20 | 20 | 12 |
| | Tremor 10-15Hz | 20 | 20 | 20 | 20 | 20 | 20 | 19 | 11 | 21 | 21 | 21 | 20 | 6 | - | - | - | 20 | 20 | 20 | 20 | 20 | 20 | 20 | 12 |
| | Tremor 15-20Hz | 20 | 20 | 20 | 20 | 20 | 20 | 19 | 11 | 21 | 21 | 21 | 20 | 6 | - | - | - | 20 | 20 | 20 | 20 | 20 | 20 | 20 | 12 |
| | Tremor 20-25Hz | 20 | 20 | 20 | 20 | 17 | 20 | 19 | 11 | 21 | 21 | 21 | 20 | 6 | - | - | - | 20 | 20 | 20 | 20 | 20 | 20 | 20 | 12 |
